## Supplementary material for "A Machine Learning Framework to Identify the Correlates of Disease Severity in Acute Arbovirus Infection": Table S1

**Table 1.** **Animals and study design**.

*Main Studies*

|  | **Groups** | | |
| --- | --- | --- | --- |
| **G1** | G1-BTV-1-2006 | G1-BTV-1-2013 | G1-control |
| *Length of study scheduled* | 21 dpi | 21 dpi | 21 dpi |
| *Postmortem* | 7-9 dpi | 21 dpi | 21 dpi |
| *Animals per group (n)* | 7 | 7 | 7 |
| *Sex/breed* | Male/Sarde | Male/Sarde | Male/Sarde |
| *Location (city, country)* | Teramo, Italy | Teramo, Italy | Teramo, Italy |

|  | **Groups** | | | |
| --- | --- | --- | --- | --- |
| **G2** | G2-BTV-1-2006 | G2-BTV-1-2013 | G2-BTV-8 | G2-control |
| *Length of study scheduled* | 7 dpi | 7 dpi | 7 dpi | 7 dpi |
| *Postmortem* | 7 dpi | 7 dpi | 7 dpi | 7 dpi |
| *Animals per group (n)* | 7 | 7 | 7 | 7 |
| *Sex/breed* | Female/Sarde | Female/Sarde | Female/Sarde | Female/Sarde |
| *Location (city, country)* | Sassari, Italy | Sassari, Italy | Sassari, Italy | Sassari, Italy |

*Additional groups*

|  | **Virus** | | |
| --- | --- | --- | --- |
|  | BTV-1_IT2006_ | BTV-1_IT2006_ | Mock |
| *Length of study scheduled* | 2 dpi | 7 dpi | 2 dpi |
| *postmortem* | 2 dpi | 7 dpi | 2 dpi |
| *Animals per group (n)* | 4 | 3 | 3 |
| *Sex/breed* | Female/Sarde | Female/Sarde | Female/Sarde |
| *Location (city, country)* | Sassari, Italy | Sassari, Italy | Sassari, Italy |
